# Endothelial cell-specific DNA methylation alterations identified in breast cancer with epigenetic deconvolution

**DOI:** 10.1101/2025.07.25.666900

**Authors:** Barbara Karakyriakou, Ze Zhang, Hanxu Lu, Lucas A. Salas, Brock C. Christensen

## Abstract

DNA methylation alterations are epigenetic modifications that regulate gene expression and contribute to carcinogenesis. Technical challenges and high costs limit the use of single-cell sequencing approaches to measure and understand cell-specific DNA methylation alterations in the tumor microenvironment (TME). Using cell type deconvolution (HiTIMED) and an interaction testing framework (CellDMC), we identified DNA methylation alterations in tumor endothelial cells (TECs) from measures in bulk biospecimens. Alterations in TECs contribute to abnormal angiogenesis, resulting in disorganized, leaky blood vessels that can hinder drug delivery and immune access. We hypothesize that TECs in the TME exhibit distinct DNA methylation patterns compared to normal endothelial cells, driving phenotypic changes that support tumor growth and vascular dysfunction. We used genome-scale DNA methylation data and a testing and validation set design (tumor, n=1,080; nontumor, n=415) to identify TEC-specific DNA methylation alterations in breast cancer. Between cohorts, we validated 4,562 TEC-specific CpGs (4,163 hypermethylated and 399 hypomethylated), and found that many map to genes regulating angiogenesis and endothelial function. Integration with mRNA data from tumor samples in both discovery and validation sets revealed genes whose expression correlates with TEC-specific DNA methylation patterns and endothelial cell proportions, highlighting potentially novel therapeutic targets. This study demonstrates the utility of cellular deconvolution to uncover cell lineage-specific epigenetic changes in cancer using bulk specimen, and identifies TEC-specific methylation alterations as novel potential targets for therapeutic intervention.

## Introduction

DNA methylation, a key epigenetic modification, is crucial in regulating gene expression and maintaining cellular identity. The addition of methyl groups to cytosine residues within gene promoter regions can lead to gene silencing^1^. DNA methylation alterations implicated in carcinogenesis and disease progression include hypomethylation of repetitive elements and hypermethylation of tumor suppressor gene promoter CpG islands^2^. However, much of the focus in cancer epigenetics has been on cancer cells themselves, in part because DNA methylation is typically measured in bulk biospecimens, leaving a gap in understanding DNA methylation alterations specific to non-tumor cells within the tumor microenvironment (TME), including endothelial cells.

The TME is a complex and dynamic ecosystem composed of various cell types that significantly influence tumor biology^3^. Among these, tumor endothelial cells (TECs) play a critical role in tumor progression due to their involvement in tumor angiogenesis—the formation of new blood vessels from existing ones that supply the tumor with nutrients and facilitate metastatic spread^4,5^. Without blood vessels, tumors cannot grow beyond a microscopic size, and metastasis, the deadliest step in the tumor progression cascade, critically depends on bidirectional tumor-vessel interactions^6^.

TECs exhibit phenotypic and functional differences from normal endothelial cells, driven, in part, by epigenetic alterations such as DNA methylation^7–9^. Investigating TEC-specific DNA methylation patterns is essential, as these changes may underlie the unique properties of TECs and their role in promoting a pro-tumorigenic environment. The endothelial cell layer is now widely recognized as a highly dynamic interface that exerts instructive gatekeeper functions on its surrounding microenvironment^10^. Breast cancer, one of the most prevalent cancer types globally for women, presents a valuable model for studying these alterations. The breast TME is highly heterogeneous and includes endothelial cells that play distinct roles in tumor progression^11^. Understanding cell-specific epigenetic changes has implications not only for breast cancer but also for other cancer types, potentially revealing novel therapeutic targets to disrupt the supportive role of the TME in cancer progression.

Identifying DNA methylation changes specific to TECs poses methodological challenges, as traditional bulk analyses can be confounded by the heterogeneous cell composition of the TME. Variations in cell proportions can obscure true cell-type-specific methylation alterations, potentially leading to misleading conclusions. However, single-cell sequencing approaches for DNA methylation are technically challenging, and very high costs limit the sample sizes needed to generalize findings for translational applications. To overcome these limitations, we use methylation and other molecular data measured in bulk tumor and normal tissue specimens. By applying cell-type deconvolution using the hierarchical tumor immune microenvironment epigenetic deconvolution algorithm, HiTIMED^12^ (v0.99.3), and CellDMC^13^ (EpiDISH v2.20.1) for interaction testing, we identified TEC-specific differential methylation in the TME. Together, HiTIMED and CellDMC provide a robust framework for identifying and investigating cell-specific molecular alterations in typical bulk biospecimens.

## Results

### Cell-type composition in breast tumor and normal tissues

The study scheme showing sample sizes and our analytic approach using Illumina 450K and EPIC methylation array data from breast cancer and normal breast tissue samples is shown in **Figure 1**. We analyzed genome-wide DNA methylation profiles from 590 breast tumors and 333 normal breast tissue samples in the discovery set first using the HiTIMED^12^ deconvolution algorithm to estimate cell-type proportions in the tumor microenvironment (TME). The results of TME deconvolution into six key cell lineages: tumor, endothelial, epithelial, stromal, myeloid, and lymphocyte are shown in **Figure 2A**. As expected, cell-type compositions differed markedly between cancer and normal tissues (**Figure 2B**). Tumor sample compositions were dominated by tumor cells, whereas normal samples displayed higher proportions of epithelial, stromal, and endothelial cells. Mean lymphocyte and myeloid cell proportions were low, with minimal variation across samples. HiTIMED^12^ results for samples in the validation set are shown in supplementary data (**Figure S1A,B)**.

**Figure 1.**
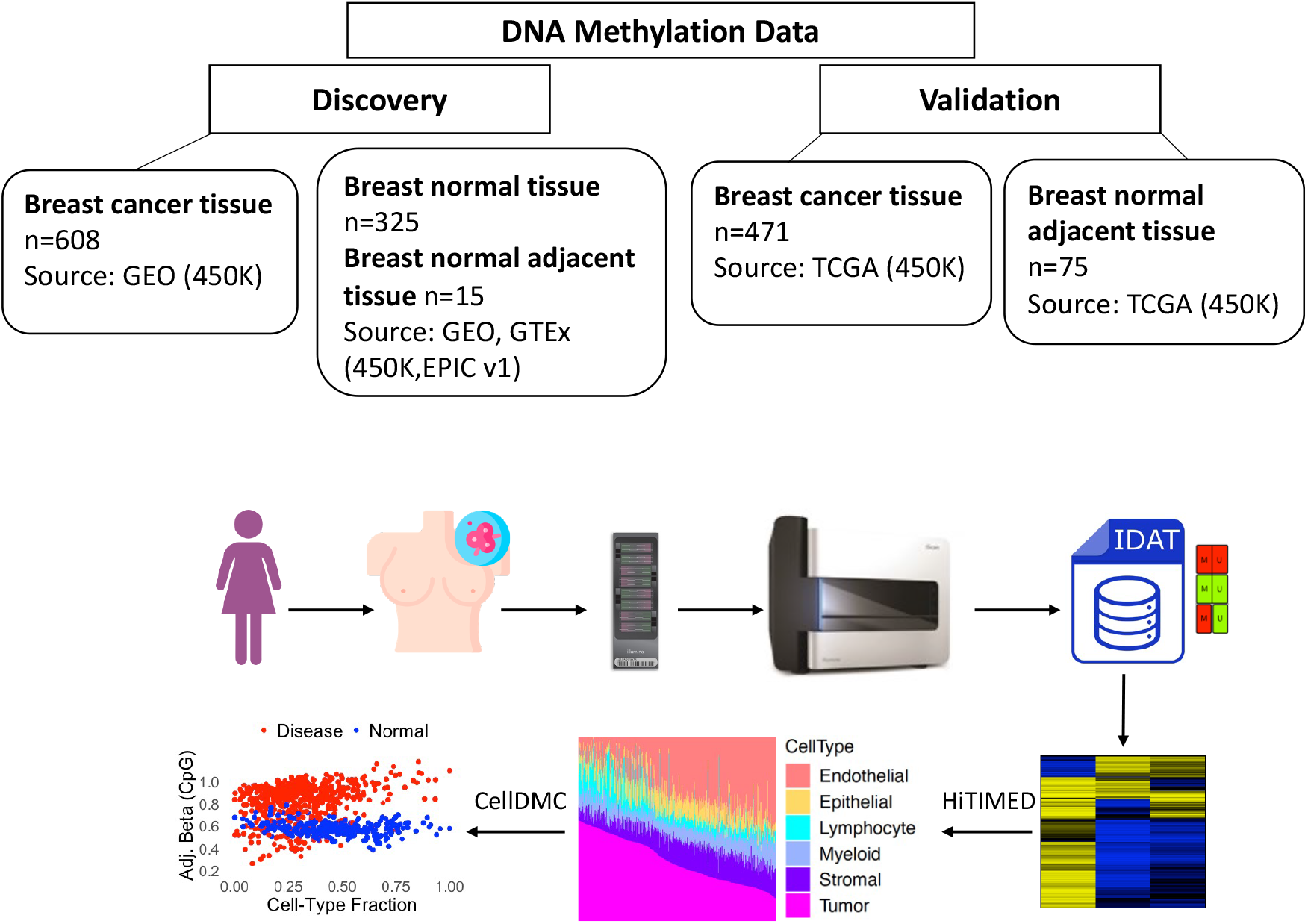
Overview of study design with analysis steps and included cohorts. Breast cancer and normal breast tissue samples were obtained and profiled using Illumina 450K and EPIC methylation arrays. The discovery set included breast cancer tissues (n=609, GEO), normal breast tissues (n=325, GEO and GTEx), and adjacent normal tissues (n=15, TCGA). The validation set comprised breast cancer (n=471) and adjacent normal (n=75) tissues from TCGA. Raw IDAT files containing red/green fluorescence signal intensities were processed to generate a beta value matrix: ratio of methylated probe intensity to the sum of methylated and unmethylated probe intensities at each CpG site, ranging from 0 (unmethylated) to 1 (fully methylated). Cell type proportions were estimated using the HiTIMED deconvolution algorithm, leveraging reference-based methylation signatures for six cell types: endothelial, epithelial, lymphocyte, myeloid, stromal, and tumor cells. The CellDMC model was applied to identify differential methylation adjusted for cell-type heterogeneity.

**Figure 2.**
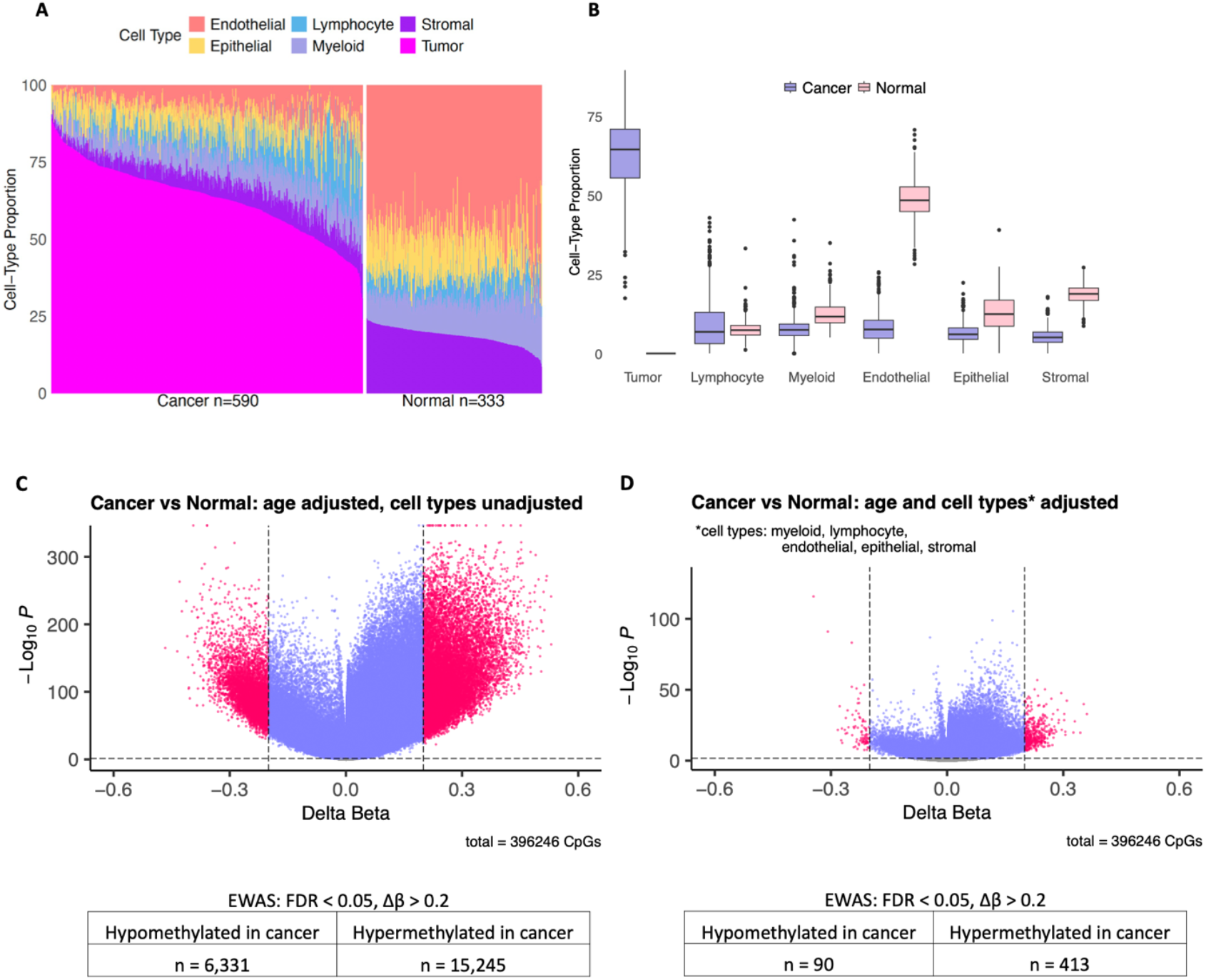
Cell-type composition and impact of cell-type adjustment on epigenome-wide differential methylation analysis in breast cancer and normal breast tissue. **A.** Stacked bar plots representing cell type proportions in discovery samples (breast cancer n=590, normal breast n=333) ranked by tumor percentage in cancer tissue and stromal percentage in normal tissue. **B.** Cell-type composition differences between breast cancer and normal breast tissue samples. Boxplots show the estimated proportions of six major cell types— Tumor, Lymphocyte, Myeloid, Endothelial, Epithelial, and Stromal—in cancer (purple) and normal (pink) samples. **C.** Volcano plot displaying differential DNA methylation in epigenome-wide association study (EWAS) between cancer and normal breast tissue samples, adjusted for age but not for cell-type composition. Each point represents a CpG site (n = 396,246). Red points indicate CpGs with both statistically significant and biologically meaningful methylation differences (|Δβ| > 0.2 and FDR-adjusted P < 0.05). **D.** Volcano plot displaying differential DNA methylation between in EWAS between cancer and normal breast tissue samples (n = 396,246 CpGs), adjusted for age and five major cell types (myeloid, lymphocyte, endothelial, epithelial, and stromal). Red points highlight CpGs with both statistically significant and biologically meaningful differences (|Δβ| > 0.2 and FDR-adjusted P < 0.05). Summary tables below each plot report the number of hypermethylated and hypomethylated CpGs identified after adjustment.

First, an epigenome-wide association study (EWAS) approach was used to compare CpG-specific DNA methylation in tumors with normal breast tissues in the discovery population. The initial EWAS included adjustment for age and identified 21,576 CpGs with significant differential methylation in tumors compared with normal tissue (FDR<0.05 and |Δβ|>0.2, **Figure 1C**). When adjusting for age and 5 cell-type proportions (myeloid, lymphocyte, endothelial, epithelial, and stromal), the scope of observed significantly differentially methylated CpG sites was dramatically reduced (503 CpGs FDR<0.05 and |Δβ|>0.2, **Figure 1D**). To support valid cell-type-adjusted modeling, we assessed multicollinearity among estimated cell-type fractions using variance inflation factor (VIF) analysis (Table S2), confirming acceptable levels for inclusion in regression models.

### Reproducible tumor endothelial cell-specific DNA methylation alterations exhibit a distinct epigenetic landscape in the breast tumor microenvironment

To identify tumor endothelial cell (TEC)-specific DNA methylation alterations, we applied the CellDMC algorithm to both discovery and validation datasets. To avoid structural zeros in CellDMC analyses, cell-type proportions were re-established using HiTIMED^12^ by setting the tissue type to “tumor” for all samples, generating low but non-zero tumor cell fractions in normal tissues. A visualization of the resulting tumor cell proportions in normal samples for both discovery and validation sets is provided in supplementary data (**Figure S2A,B**). To ensure sufficient statistical power for the validation analysis, the validation set included the normal samples from the discovery cohort.

In the discovery set, out of 396,246 CpG sites tested, 25,164 hypermethylated, and 1,528 hypomethylated TEC-specific CpGs were identified as significantly differentially methylated (FDR ≤ 0.05, and |coef.| >0.0062 corresponding to ≥5% methylation change in endothelial cells) (**Figure 3A)**.To ensure specificity, we removed CpGs that were also differentially methylated in other cell types in the same direction. Specifically, 12,949 hyper- and 524 hypomethylated CpGs overlapped with tumor cell signals; 2,143 hyper- and 13 hypomethylated CpGs overlapped with epithelial signals; and additional overlaps were observed with stromal and myeloid compartments. After filtering, we identified 10,041 hypermethylated and 990 hypomethylated CpGs uniquely altered in TECs (**Figure 3B**).

**Figure 3.**
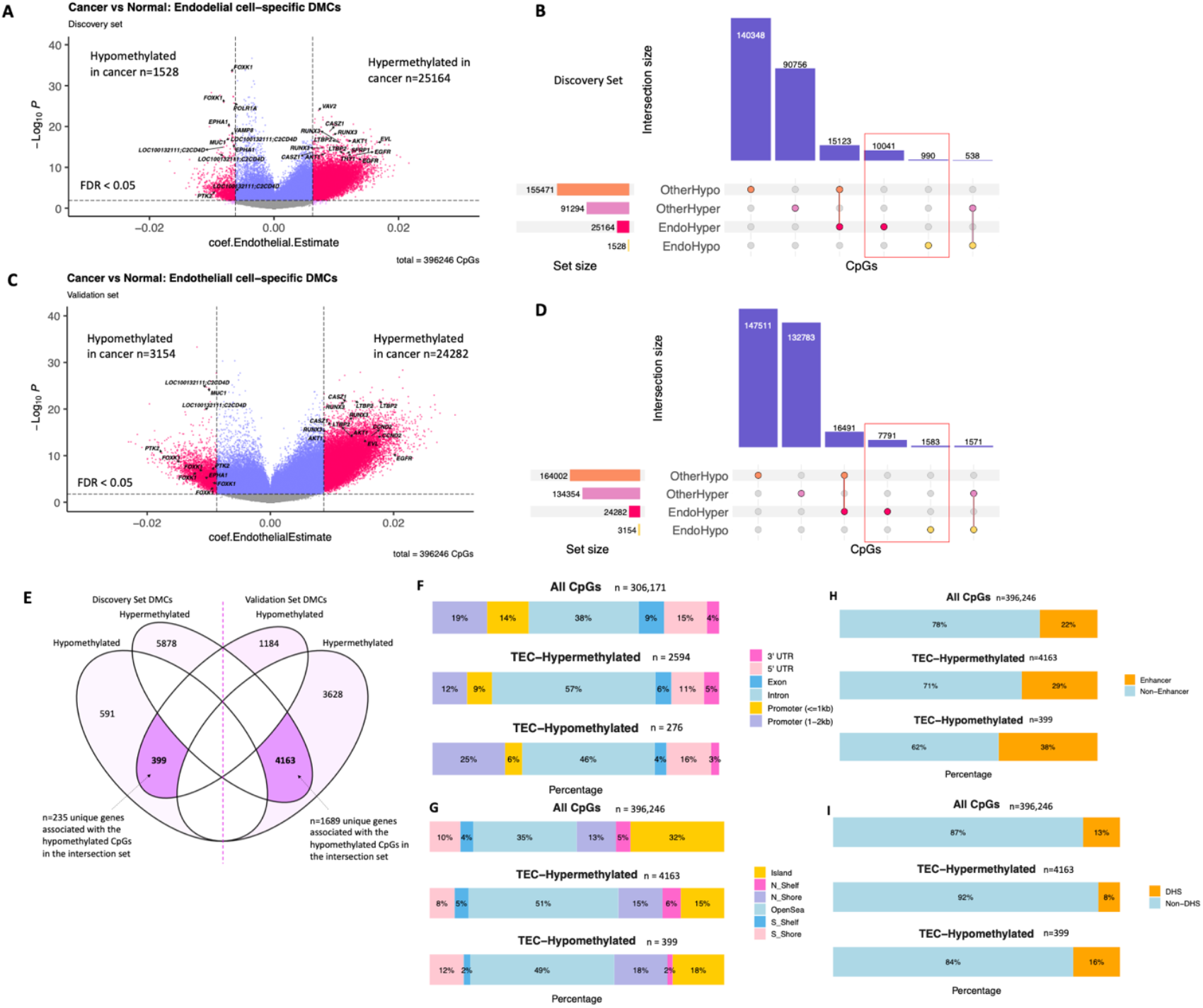
Identification, validation, and functional annotation of tumor endothelial cell (TEC)-specific differentially methylated CpGs (DMCs) in breast cancer. **A.** Volcano plot of TEC-specific DMCs in the discovery set. Out of 396246 total CpGs, 99058 were identified as differentially methylated in tumor endothelial cells (FDR <= 0.05) after adjusting for age. **B.** Upset plot showing uniqueTEC-specific DMCs in the discovery set (10041 hypermethylated and 990 hypomethylated) after filtering out CpGs that are DM in the same direction with other cell-types. **C.** Volcano plot of TEC-specific DMCs in the validation set. Out of 396246 total CpGs, 143463 were identified as TEC-specific DMCs (FDR <= 0.05) after adjusting for age. **D.** Upset plot showing uniqueTEC-specific DMCs in the validation set (7791 hypermethylated and 1593 hypomethylated) after filtering out CpGs that are DM in the same direction with other cell-types. **E.** The intersection of TEC-specific DMCs across both sets includes 4,562 CpGs (4,163 hypermethylated and 399 hypomethylated) which are associated with a total of 1,924 genes. **F.** Stacked bar plots showing the percentage of CpGs in different genomic regions for all tested CpGs and for TEC-specific hypermethylated and hypomethylated CpGs. **G.** Stacked bar plots showing distribution of all tested CpGs and TEC-specific DMCs relative to CpG islands. **H.** Regulatory distribution of TEC-specific DMC compared to all CpGs in enhancer regions. **I.** Regulatory distribution of TEC-specific DMC compared to all CpGs in DNase I hypersensitive sites.

In the validation set, the CellDMC model identified 24,282 hypermethylated and 3,154 hypomethylated TEC-specific DMCs (FDR ≤ 0.05, |coef.| >0.0087 corresponding to ≥5% methylation change) (**Figure 3C**). Using the same filtering strategy, we excluded 14,736 hyper- and 1,553 hypomethylated CpGs that overlapped with tumor cell signals, along with additional overlaps from epithelial, stromal, and myeloid cells. This resulted in 7,791 hypermethylated and 1,593 hypomethylated uniquely TEC-specific CpGs (**Figure 3D**). To assess reproducibility, we intersected the uniquely TEC-specific DMCs from both datasets. A total of 4,562 CpGs were consistently differentially methylated across discovery and validation sets, including 4,163 hypermethylated and 399 hypomethylated sites (**Figure 3E**), highlighting a robust and reproducible TEC-specific epigenetic signature in breast cancer. These CpGs mapped to 1,689 unique genes for the hypermethylated set and 235 genes for the hypomethylated set, suggesting both widespread and targeted transcriptional regulatory potential.

To further characterize the biological relevance of these alterations and assess broader genomic patterns, we analyzed the distribution of hyper- and hypomethylated TEC-specific CpGs across annotated gene regions (**Figure 3F**). Compared to all tested CpGs associated to genes (n = 306,171), TEC-specific hypermethylated CpGs were more frequently located within gene bodies, particularly in introns. In contrast, hypomethylated CpGs in TECs were disproportionately enriched in promoter regions upstream of transcription start sites (TSS1500). Analysis of the CpG island context revealed an enrichment of both hyper- and hypomethylated CpGs in open sea regions (**Figure 3G**). TEC-specific hypermethylated CpGs showed increased enhancer localization compared to the background set, with a similar trend observed for hypomethylated CpGs (**Figure 3H**). TEC-specific hypomethylated CpGs were enriched in DNase I hypersensitive sites (DHS), whereas hypermethylated CpGs were depleted in DHS (**Figure 3I**).

### TEC-specific DMCs map to angiogenesis genes

We further examined the gene associations of tumor endothelial cell (TEC)-specific differentially methylated CpGs (DMCs) identified in the intersected discovery and validation sets. Notably, among the uniquely differentially methylated CpGs identified in both the discovery and validation sets, several mapped to *CASZ1, EGFR, EVL, RUNX3, SFRP1*, and *THY1*, genes known to regulate angiogenesis and endothelial cell biology. Across both datasets, we identified DMCs associated with genes in the *VEGF* signaling pathway: *AKT1, AKT2, CDC42, CHP2, MAPK13, PIK3CD, PIK3R2, PIK3R5, PLA2G10, PLA2G4A, PRKCA, PRKCB, PRKD1, PTK2, SPHK2* and *VEGFA*, and hallmark angiogenesis genes including *CCND2, ITGAV, LRPAP1, MSX1, POSTN, PTK2, VAV2*, and *VTN*. In the intersected set of high-confidence TEC-specific DMCs, nearly all of these angiogenesis-associated genes were retained. This finding reinforces the biological relevance of the intersected CpGs and their potential as robust biomarkers or therapeutic targets in tumor vascular remodeling.

Representative examples of TEC-specific DMCs mapped to angiogenesis-related genes are shown for *RUNX3* (**Figure 4A**), *EGFR* (**Figure 4B**), and *CASZ1* (**Figure 4C**). These genes showed clusters of hypermethylated CpGs in TECs across multiple gene isoforms. Heatmaps reveal a consistent methylation shift between normal and cancer tissues in both discovery and validation sets, with normal samples exhibiting lower methylation and tumor samples showing increased methylation, indicating TEC-specific epigenetic remodeling within these angiogenic regulators.

**Figure 4.**
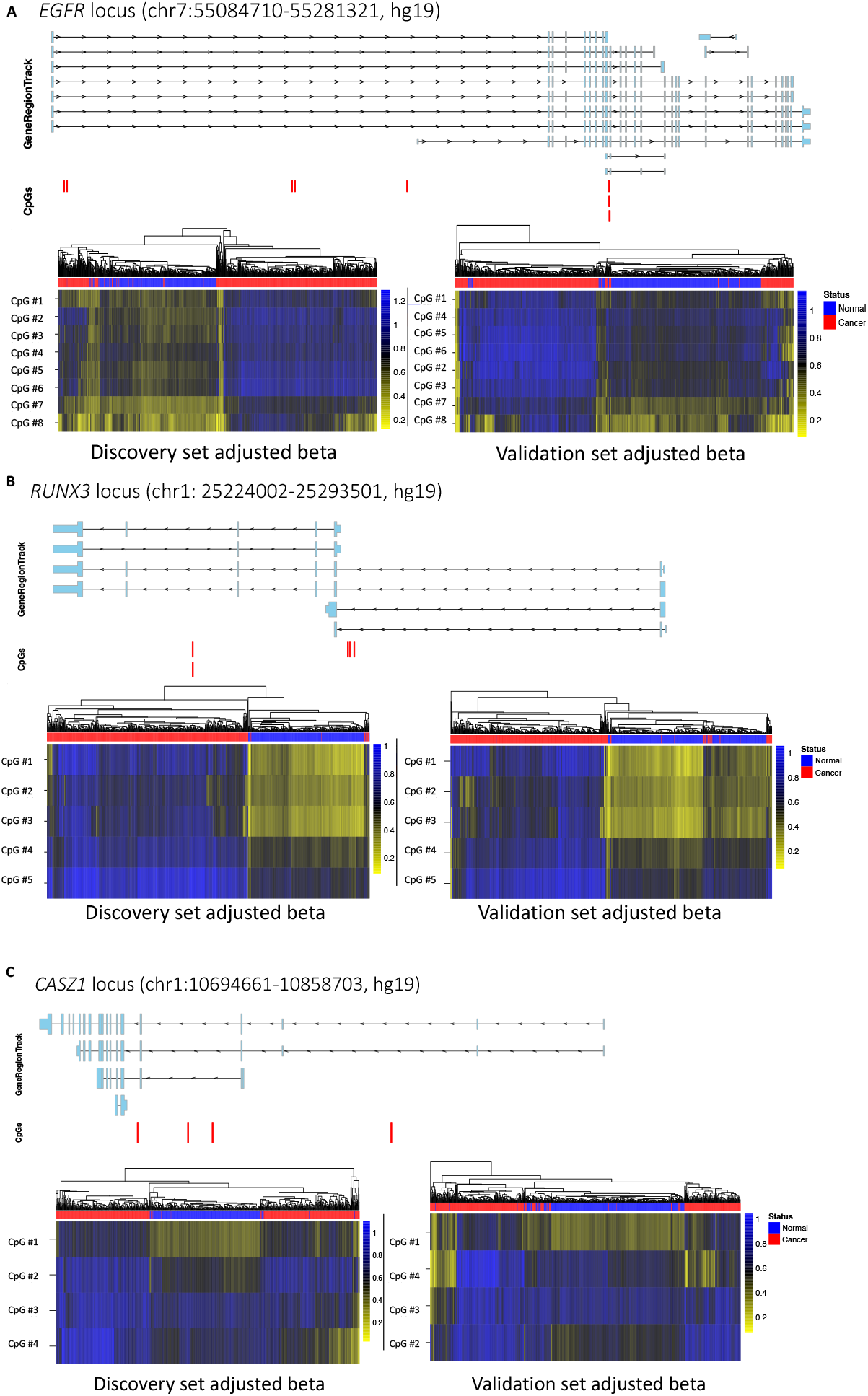
TEC-specific differentially methylated CpGs in angiogenesis-related genes in breast cancer. **A.** Genomic view and Manhattan distance heatmaps of CpG methylation in the *EGFR* locus, showing eight TEC-specific hypermethylated CpGs in tumor versus normal samples within the discovery and validation sets. **B.** Genomic view and Manhattan distance heatmaps of five TEC-specific hypermethylated CpGs at the *RUNX3* locus. **C.** Genomic view and Manhattan distance heatmaps of four TEC-specific hypermethylated CpGs at the *CASZ1* gene locus. Red vertical bars indicate the exact CpG positions within each gene region.

### Endothelial cell proportion correlates with the expression of genes linked to TEC-Specific DMCs

To investigate the transcriptional relevance of tumor endothelial cell (TEC)-specific DNA methylation alterations, we analyzed gene expression data from 245 breast cancer samples in the discovery set with matched Agilent microarray mRNA profiles. We focused on genes associated with CpGs in the intersected TEC-specific differentially methylated (DMC) set and assessed their correlation with endothelial cell proportions across samples. The top correlated gene, *LHFP*, exhibited a strong positive correlation between expression and endothelial content (r = 0.6, p < 0.001). Overall, 13 genes demonstrated moderate to strong correlation (r ≥ 0.5), as shown in **Table 1**. Many of the CpGs associated with these genes were hypermethylated and mapped to gene body and enhancer regions, suggesting epigenetic regulation through non-promoter elements (**Table 2**).

**Table 1.** Top genes showing consistent expression-endothelial cell proportion correlation in discovery and validation datasets.

| Gene | Pearson's correlation coefficient |  | Adjusted P-value (BH) |  |
| --- | --- | --- | --- | --- |
|  | Discovery | Validation | Discovery | Validation |
| *LHFP* | 0.60 | 0.70 | 1.18E-21 | 2.39E-67 |
| *NDN* | 0.56 | 0.64 | 4.80E-19 | 2.34E-52 |
| *NID1* | 0.56 | 0.51 | 4.80E-19 | 1.04E-30 |
| *MSRB3* | 0.56 | 0.58 | 7.40E-19 | 2.89E-41 |
| *GPR124* | 0.55 | 0.60 | 2.77E-18 | 1.65E-45 |
| *DLC1* | 0.55 | 0.50 | 2.77E-18 | 6.23E-30 |
| *RHOJ* | 0.55 | 0.66 | 4.20E-18 | 5.45E-57 |
| *JAM2* | 0.53 | 0.62 | 3.70E-17 | 6.77E-48 |
| *LOC399959* | 0.53 | 0.48 | 3.96E-17 | 2.21E-27 |
| *RUNX1T1* | 0.53 | 0.44 | 5.85E-17 | 8.73E-23 |
| *RBMS3* | 0.53 | 0.57 | 1.49E-16 | 2.47E-40 |
| *LTBP2* | 0.52 | 0.52 | 2.78E-16 | 9.06E-32 |
| *HTRA1* | 0.50 | 0.59 | 6.58E-15 | 8.89E-43 |

**Table 2.** Annotation of TEC-specific DMCs associated with genes correlated to endothelial cell proportion.

| Gene name | CpG | Relation to Island | Genic Group | Enhancer |
| --- | --- | --- | --- | --- |
| **Hypermethylated in TECs** |  |  |  |  |
| *LHFP* | cg05099186 | OpenSea | Body | TRUE |
| *NDN* | cg18552939 | S_Shore | TSS200 |  |
| *MSRB3* | cg05475386 | OpenSea | Body | TRUE |
| *GPR124* | cg14922775 | OpenSea | Body |  |
|  | cg06009497 | N_Shelf | Body |  |
| *DLC1* | cg01070197 | OpenSea | Body |  |
|  | cg22732636 | OpenSea | TSS1500 |  |
| *RHOJ* | cg07157030 | OpenSea | 5'UTR;1stExon | TRUE |
| *LOC399959* | cg05005343 | OpenSea | Body | TRUE |
|  | cg16865908 | OpenSea | Body | TRUE |
|  | cg24213115 | OpenSea | Body | TRUE |
|  | cg07281370 | OpenSea | Body | TRUE |
|  | cg06749053 | OpenSea | Body | TRUE |
|  | cg05678526 | OpenSea | Body |  |
|  | cg26363555 | OpenSea | Body |  |
|  | cg17389813 | OpenSea | Body |  |
|  | cg03393223 | OpenSea | TSS1500 | TRUE |
| *RUNX1T1* | cg07538339 | OpenSea | 5'UTR;1stExon |  |
| *JAM2* | cg10466421 | N_Shore | TSS1500 |  |
| *RBMS3* | cg22494026 | OpenSea | Body | TRUE |
| *LTBP2* | cg14358448 | OpenSea | Body | TRUE |
|  | cg16056219 | OpenSea | Body | TRUE |
|  | cg26011692 | Island | Body |  |
|  | cg16239257 | Island | Body |  |
|  | cg14986500 | Island | Body |  |
|  | cg11718443 | Island | Body |  |
|  | cg14340889 | OpenSea | Body | TRUE |
| *HTRA1* | cg03974204 | OpenSea | Body |  |
| **Hypomethylated in TECs** |  |  |  |  |
| *NID1* | cg07157830 | Island | TSS1500 |  |
|  | cg04837025 | Island | TSS1500 |  |

To validate these findings, we repeated the analysis using RNA-sequencing data from the independent validation cohort. Consistent with the discovery set, *LHFP* emerged as the top-correlated gene (**Figure 5A**), and the core set of 13 genes was consistently ranked among the top correlations in both datasets despite differences in experimental platforms. This cross-platform concordance strengthens the evidence for a biologically meaningful association between TEC-specific methylation and transcriptional activation in endothelial cells.

**Figure 5.**
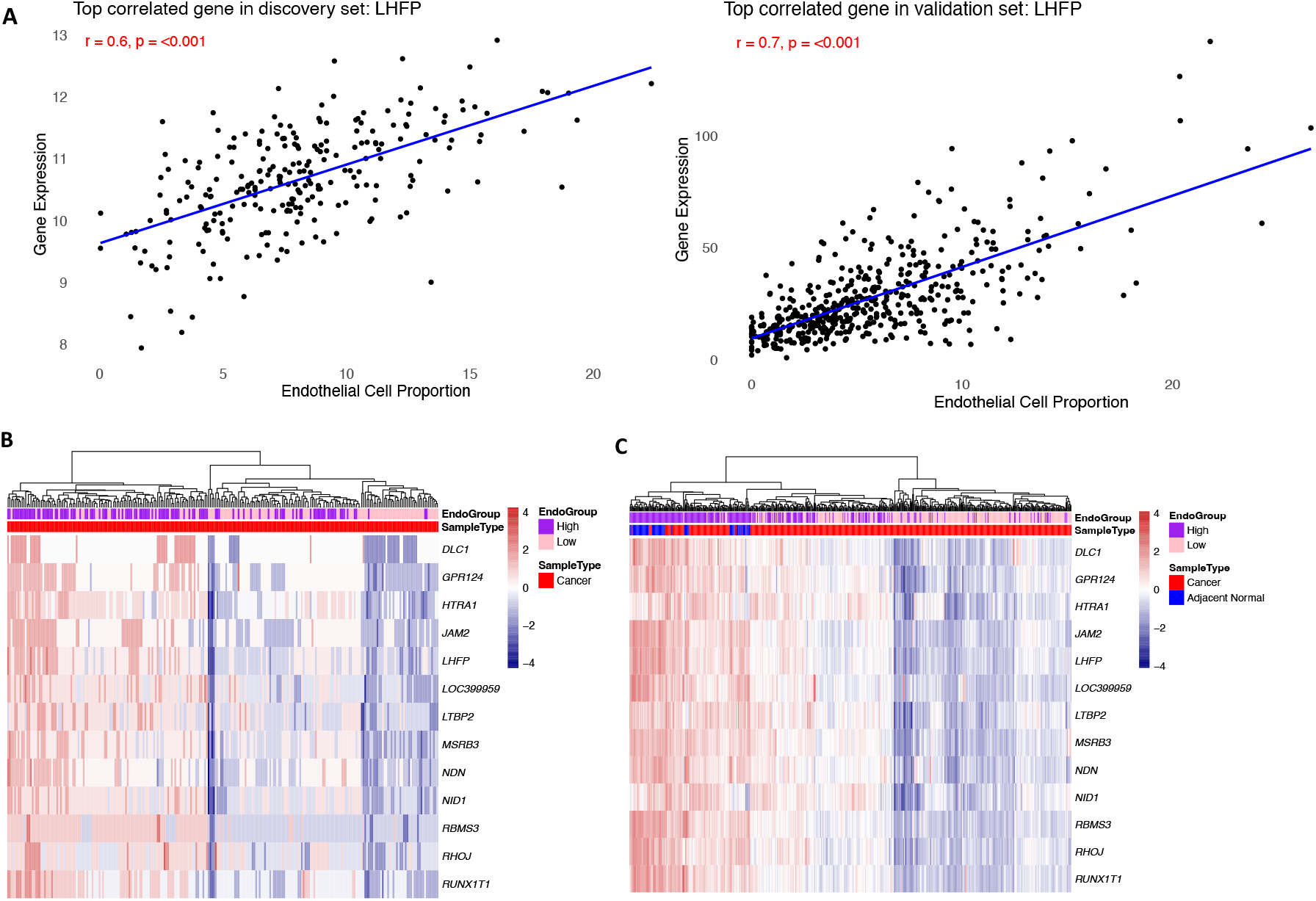
Correlation between endothelial cell proportion and gene expression for genes associated with TEC-specific DMCs in breast tumors. **A.** Scatter plots showing strong positive correlation between *LHFP* expression and endothelial cell proportion in the discovery set (r = 0.6, p < 0.001) and validation set (r = 0.7, p < 0.001). **B.** Euclidian distance heatmap of top correlated genes in the discovery set (Agilent microarray gene expression data, n=245). Samples are annotated as low or high endothelial group based on the median endothelial cell proportion. **C.** Euclidian distance heatmap of top correlated genes in the validation set (RNA-seq TPM, breast cancer n=469 and adjacent normal n=64). Samples are annotated by tissue type (cancer or adjacent normal) and endothelial group (low or high).

To further characterize the transcriptional pattern of these genes, we visualized their expression in both datasets using heatmaps. In the discovery set (Agilent microarray expression data; n = 245), samples were stratified by endothelial cell proportions into high and low groups (**Figure 5B**). In the validation set (RNA-seq TPM data; n = 533), samples were annotated by tissue type (cancer or adjacent normal) and endothelial group (**Figure 5C**). In both cohorts, adjacent normal tissues and tumors with high endothelial content exhibited elevated expression of the TEC-associated gene set compared to low-endothelial tumors.

As a negative control, we selected a random gene set of the same size as the TEC-specific DMC-associated gene set and evaluated their expression correlation with endothelial cell proportions. In both the discovery and validation datasets, the correlation coefficients for the random gene sets were consistently lower compared to those of the TEC-associated gene set, further supporting the specificity of the observed associations. Additionally, we assessed whether the TEC-specific gene set was correlated with the proportions of other cell types within the tumor microenvironment. Across all other cell types, the expression of these genes showed no significant correlation except for stromal cells (supplementary data).

## Discussion

Our study underscores the importance of and opportunity to identify cell-type-specific epigenetic alterations in the TME using DNA methylation data measured in bulk biospecimens. By identifying differentially methylated CpGs (DMCs) specific to tumor endothelial cells (TECs), we provide novel insights into the unique epigenetic regulation of genes involved in angiogenesis and other pathways critical to tumor progression. The specificity of TEC-associated DMCs highlights the limitations of bulk tissue analyses, which obscure the nuanced methylation landscapes of individual cell types. For instance, we observed that many TEC-specific DMCs were either non-significant or showed opposite methylation directionality in other cell types, emphasizing the distinct epigenetic identity of TECs. These findings validate the robustness of our deconvolution-based approach and illustrate the critical need to account for cell-type-specific effects when interrogating the TME.

Functional annotation of TEC-specific methylation alterations revealed divergent regulatory potential. When examining the genomic context of TEC-specific DMCs, we found distinct patterns of localization between hyper- and hypomethylated CpGs. Hypermethylated sites were enriched in gene bodies and enhancer regions, whereas hypomethylated CpGs were predominantly located in promoter regions. Analysis of the CpG island context further showed that both hypermethylated and hypomethylated CpGs were enriched in open sea regions, which often harbor distal regulatory elements such as enhancers. Notably, TEC-specific hypomethylated CpGs were also enriched in DHS, suggesting that these alterations preferentially occur in regions of accessible chromatin associated with active regulatory function. This contrasts with the depletion of hypermethylated CpGs in DHS, implying that hypermethylation may target regions that are normally inaccessible in bulk tissue but become selectively remodeled in TECs. These findings support a model in which the TEC methylome is remodeled to shape cell-specific regulatory landscapes: hypomethylation may facilitate the activation of pre-accessible regulatory elements, while gene body hypermethylation may contribute to modulation of gene expression, potentially influencing transcriptional elongation or alternative splicing. Although promoter DNA methylation is well known to suppress gene expression, gene body DNA methylation can regulate cell-type-specific alternative promoters and paradoxically enhance transcription elongation efficiency^14^. Together, these patterns illustrate the distinct epigenetic mechanisms by which TECs acquire a tumor-supportive regulatory phenotype.

TEC-specific DMCs mapped to thousands of genes, with several loci showing dense clustering of hypermethylated sites— with the largest cluster comprising 108 TEC-specific hypermethylated CpGs associated with DAXX, a gene implicated in chromatin remodeling and cell fate regulation that is also known to contribute to tumorigenesis^15–17^. Notably, we identified clusters of TEC-specific DMCs within several genes with established roles in angiogenesis, highlighting coordinated epigenetic alterations at these loci. For example, *EGFR* harbored eight hypermethylated CpGs within its gene body, some of which were also associated with enhancer regions. *EGFR* facilitates crosstalk with VEGF signaling pathways in the TME, promoting angiogenesis and supporting tumor progression^18–21^. *RUNX3* contained five hypermethylated CpGs, and *CASZ1* had four hypermethylated CpGs within gene body regions, with some overlapping enhancers. *RUNX3*, a known tumor suppressor, inhibits angiogenesis by binding the *VEGF* promoter and downregulating VEGF production^22–24^, while *CASZ1* is a transcription factor essential for angiogenic sprouting and vessel morphogenesis^25–27^.

We also identified a cluster of five hypermethylated CpGs within the body of *PRKCA*, a key member of the human VEGF signaling pathway^28–30^, known to promote angiogenic activity of endothelial cells via induction of *VEGF*^31^. More broadly, multiple TEC-specific DMCs were consistently associated with genes in the VEGF signaling pathway and with genes from the Hallmark Angiogenesis gene set^32^, including *AKT1*, a core component of the PI3K–AKT pathway, which promotes pathological angiogenesis by modulating vascular permeability, endothelial cell activation, and vessel maturation^33–40^.

In addition, TEC-specific hypermethylated CpGs were detected in *EVL* and *THY1*, genes implicated in endothelial function and vascular remodeling. *EVL* (Ena-VASP-like) regulates actin cytoskeleton dynamics and endothelial sprouting behavior^41,42^, while *THY1* (CD90) mediates endothelial adhesion and migration—both essential processes for neovascularization^43–45^.

Together, these findings illustrate the functional relevance of TEC-specific DMCs and their potential contribution to a pro-angiogenic and pro-tumorigenic microenvironment. Collectively, our results highlight the enrichment of endothelial cell-specific CpG alterations in genes associated with angiogenesis and further reveal consistent, biologically meaningful TEC-specific DNA methylation remodeling in the breast tumor microenvironment, nominating high-confidence candidate loci for functional validation and potential therapeutic targeting.

The observation of CpGs with inverse methylation patterns between TECs and other cell types suggests possible intercellular regulatory mechanisms. It remains to be determined whether these patterns result from direct cell-cell interactions, microenvironmental signaling, or differential responses to stressors such as hypoxia or VEGF. Future multi-omic studies integrating methylation, transcriptomic, and proteomic data will be essential to elucidate these mechanisms and their downstream biological consequences.

Importantly, the identification of TEC-specific DNA methylation signatures offers valuable insight into the epigenetic landscape of the tumor vasculature. By identifying TEC-specific methylation signatures enriched in angiogenesis- and hypoxia-related pathways, our work lays the groundwork for future functional studies and highlights potential biomarkers or therapeutic entry points to develop combination treatment approaches that impact the TME.

While direct targeting of overexpressed proteins or transcript variants remains a feasible and clinically tractable strategy, characterizing TEC-specific epigenetic alterations provides complementary opportunities—for instance, by enabling cell-type-specific biomarker discovery, elucidating regulatory mechanisms or guiding the development of epigenetic editing strategies. In particular, targeting epigenetic regulators in TECs could represent a promising approach for disrupting tumor vascularization while minimizing off-target effects on non-endothelial cell types. Emerging tools such as CRISPR-based epigenome editors—for example, dCas9 fused to DNMT3A or TET1 catalytic domains—offer locus-specific modulation of DNA methylation and may enable future functional interrogation or therapeutic targeting of TEC-specific epigenetic alterations^46^.

In addition to known angiogenic regulators, our analysis revealed novel candidate loci with strong TEC-specific methylation signals that warrant further investigation. By integrating mRNA expression data with TEC-specific methylation profiles, we identified a subset of genes whose expression strongly correlates with endothelial cell abundance in breast tumors. These findings highlight putative endothelial-specific targets, including novel candidates such as *LHFP*, that may contribute to the functional phenotype of tumor-associated endothelial cells.

Among these, *RUNX1T1*^47^, a transcription factor previously implicated in angiogenic regulation, and *RHOJ*^48–52^, previously known to play an important role in endothelial cell biology and angiogenesis, were also among the top correlated genes, supporting a potential functional link between TEC-specific methylation and endothelial transcriptional activity. In addition, *LOC399959 (also known as the MIR-100/LET-7A-2/MIR-125B-1 cluster host gene)*^53,54^, a poorly characterized long noncoding RNA that hosts a microRNA previously implicated in multiple cancer types and often associated with unfavorable prognosis, harbored multiple TEC-specific hypermethylated CpGs in open sea enhancer regions, highlighting its potential as novel regulatory element in TECs.

Together, these findings suggest possible regulatory roles in tumor endothelial cell function and nominate these loci as promising epigenetic targets for future mechanistic studies and therapeutic exploration. Consistent expression patterns identified across both discovery and validation cohorts further support the relevance of the TEC-associated gene set, with elevated expression observed in adjacent normal tissues and tumors enriched for endothelial cells. This pattern suggests that the identified gene signature reflects a conserved endothelial transcriptional program preserved across normal vasculature and more endothelial-rich tumor contexts. Tumors with higher endothelial content may retain features of a more “normal-like” vascular phenotype, potentially indicative of active angiogenesis, whereas tumors with low endothelial proportions may exhibit a dysfunctional or regressed vasculature, a phenotype often linked to poorly perfused, hypoxic microenvironments. These observations imply that certain aggressive tumors, in their drive to vascularize, may partially recapitulate aspects of normal endothelial biology—possibly as a mechanism to support growth, metabolic adaptation, and survival under stress.

Interestingly, the TEC-associated gene set identified in the mRNA-endothelial cell proportion correlation analysis showed minimal correlation with other cell types in the TME, except for stromal cells. This observation suggests that genes identified as TEC-associated may also be upregulated in stromal components, which include cancer-associated fibroblasts (CAFs) and pericytes—cell types known to contribute to tumor angiogenesis^55–63^. This overlap raises the possibility that a subset of the TEC-specific transcriptional program may be shared with stromal cells involved in vascular remodeling and angiogenic support.

Using cell deconvolution and a statistical interaction testing framework, we demonstrate the value and opportunity of high-resolution, cell-type-specific DNA methylation profiling in bulk biospecimen data for advancing our understanding of tumor-associated endothelial biology. Our findings reinforce the idea that key regulatory events may be overlooked when analyzing bulk tissue, underscoring the importance of resolving epigenetic alterations within cell types or cell lineages. Our analytical framework is versatile and can be readily extended to other stromal and immune cell populations within the TME, such as CAFs, to elucidate their distinct epigenetic contributions to tumor progression. However, large sample sizes are required to be adequately powered in the interaction testing framework. Future work will apply this framework to other solid tumor types to determine the extent of shared and distinct endothelial-specific methylation alterations across cancer types. Identification of conserved epigenetic alterations in TECs could ultimately inform the development of targeted or combination therapeutic strategies.

## Methods

### Study Population Data

Breast cancer and normal breast tissue DNA methylation data were obtained from the Gene Expression Omnibus (GEO), The Cancer Genome Atlas (TCGA), and the Genotype-Tissue Expression (GTEx) project. All samples were profiled using Illumina Infinium HumanMethylation450 BeadChip or MethylationEPIC arrays. The **discovery set** included breast tumor tissues from GSE60185 (n=239), GSE66313 (n=40), GSE84207(n=330), adjacent normal breast tissues from GSE66313 (n=15), and normal tissues from GSE60185 (n=46), GSE74214 (n = 18), GSE88883 (n = 100), GSE101961 (n = 121), and GTEx (n = 40). Samples were restricted to females, with an emphasis on capturing a broad age range (supplementary data, Table S1). The availability of matched gene expression data was also considered when possible: GSE58212 (n=285) mRNA samples matched with samples in GSE84207 methylation set. Tumor samples were from primary sites, adjacent normal samples were matched to corresponding tumors, and normal tissues were derived from mammary tissues. GEO datasets were selected based on the availability of raw IDAT files, clear tissue-type annotations, and sufficient metadata. The **validation set** comprised breast cancer (n = 471) and adjacent normal (n = 75) tissues from TCGA. TCGA samples were selected from the BRCA cohort, restricting to female samples and requiring primary tumor sites, with matched adjacent normal tissues when available, with available DNA methylation data (Table S1). Data for all TCGA projects were downloaded with the TCGAbiolinks R package (v2.32.0)^64^, setting the parameters data.category to “DNA Methylation,” platform to “Illumina Human Methylation 450” and data.type to “Raw Intensities.”

### Data Preprocessing and Quality Control

Raw intensity data (IDAT files) were processed using the minfi (v1.50.0)^65^ and ENmix (v1.40.2)^66^ toolkits, Bioconductor packages designed for comprehensive analysis of Illumina Methylation BeadChip Arrays, including quality control and normalization. Probes containing single nucleotide polymorphisms (SNPs) or those classified as cross-reactive were excluded at both the locus and sample levels.

Quality assessment of samples and probes was conducted using multiple criteria, including out-of-band detection P-values to ensure accurate signal detection, mean fluorescence intensity of bisulfite conversion control probes to evaluate methylation efficiency, and the number of beads per probe to confirm adequate probe coverage. Within-array correction for background fluorescence and dye biases was conducted using minfi’s preprocessFunnorm with the default setting method for background subtraction, Noob (normal-exponential out-of-band), followed by first two principal components of the control probes to infer the unwanted variation, ensuring uniform signal correction^67^. Following quality control and correction, beta values were calculated for each CpG site, representing the proportion of methylated to unmethylated signals. These beta values, ranging from 0 to 1, provided a continuous measure of DNA methylation intensity at each locus. Data were filtered to include only probes common to the 450K array, excluding cross-reactive probes and probes that failed quality control. The final dataset comprised high-quality methylation measurements across 396,246 CpG loci, after excluding 18 breast cancer and 7 adjacent normal breast samples from the discovery set that failed quality control.

To address batch effects arising from variations in experimental conditions and sample processing between independent datasets, we implemented robust batch correction techniques. We employed ComBat^68^, an empirical Bayes-based batch correction algorithm, to harmonize DNA methylation profiles across datasets. This approach mitigated non-biological variability, ensuring consistency and reliability in downstream analyses.

### Statistical Analysis

We performed an Epigenome-Wide Association Study (EWAS) to systematically identify DNA methylation differences across the genome associated with our phenotype of interest. EWAS enables unbiased, high-resolution detection of differentially methylated CpG sites by comparing methylation levels between groups. To control for potential confounding factors, we used the *limma*^69^ (v3.60.6) R package to adjust for age and subsequently for estimated cell-type proportions in our linear modeling framework. To ensure that the identified CpGs represent biologically meaningful differences, we applied a Δβ threshold of 0.2, which reflects an effect size corresponding to a ≥20% change in methylation. This effect size cutoff is commonly used in EWAS studies to prioritize CpGs with substantial methylation differences that are more likely to have functional relevance. Multicollinearity diagnostics and inference for fixed effects were performed using the *lmerTest* R package, which implements the Satterthwaite’s method or Kenward-Roger’s approximations for denominator degrees of freedom^70^.

To quantify cell type proportions in the TME, we applied HiTIMED^12^, a reference-based cell-type deconvolution algorithm, to batch effect-corrected beta value matrices. HiTIMED^12^ deconvolutes DNA methylation profiles into three major TME components: tumor, angiogenic, and immune, with further resolution into specific cell types. The HiTIMED algorithm is built on tumor-type-specific models for 20 carcinoma types, including breast cancer, and was developed using discovery data from 6,726 samples encompassing carcinoma and matched normal or normal-adjacent tissue. We applied HiTIMED to all normal (n = 333) and tumor (n = 590) breast tissue samples to resolve the third hierarchical layer of the algorithm for six specific TME cell types: tumor, endothelial, epithelial, stromal, myeloid, and lymphocytes. Although HiTIMED^12^ can resolve additional cell types, particularly immune cell subsets, to preserve statistical power in subsequent analyses, we restricted the scope of cell types returned.

To identify differential DNA methylation attributable to specific cell types, we employed CellDMC^13^, a validated statistical algorithm that integrates cell-type fraction data to perform cell-type-specific analyses. CellDMC analyzes DNA methylation patterns, modeling the interactions between phenotypes (e.g., tumor versus normal) and cell-type proportions to uncover differential methylation patterns specific to individual cell lineages or cell types. We used CellDMC to analyze the endothelial cell component of the TME to uncover differential DNA methylation patterns specific to tumor endothelial cells adjusting for age. Differentially methylated CpGs (DMCs) were defined using a false discovery rate (FDR) threshold of ≤ 0.05.

To further refine biologically meaningful changes, we applied an additional filter based on effect size, selecting CpGs with a minimum methylation shift of ≥5% between cancer and normal phenotypes. To estimate this 5% methylation shift, we multiplied the average endothelial cell proportion in the cancer samples by the CellDMC interaction coefficient. Specifically, we set the interaction coefficient threshold based on the average endothelial cell proportion in cancer samples. For instance, if the average endothelial fraction in cancer was ∼8%, then an interaction coefficient of ≥ 0.0062 corresponds to an average methylation shift of 0.0062 × 8 = 0.05 (i.e., 5%) in cancer samples. The actual methylation shift will vary by sample according to its endothelial cell proportion.

For visualization and interpretation, we adjusted the original beta values for CpGs identified as endothelial-specific by CellDMC. This was done by multiplying the interaction coefficient for each CpG by the endothelial cell proportion for each sample and the phenotype indicator (0 for normal, 1 for cancer), then adding this value to the original beta value. Since phenotype = 0 for normal samples, their beta values remained unchanged.

To enable the modeling of phenotype–cell-type interactions in cancer versus normal comparisons using the CellDMC^13^ framework, it is essential that all modeled cell types have non-zero proportions across both phenotypic groups. CellDMC^13^ performs interaction modeling by incorporating both phenotype (e.g., cancer vs. normal) and cell-type fraction terms; therefore, a structural zero—such as a tumor cell proportion of exactly 0% in all normal samples— precludes estimation of valid interaction coefficients. This limitation arises from the model’s design, which assumes continuous variation in cell-type proportions across conditions. When a cell type is entirely absent from one group, the model cannot estimate its contribution to phenotype-specific methylation changes, leading to loss of power and biased results.

To address this, we re-estimated cell-type proportions using HiTIMED^12^ by setting the tissue type to tumor for all samples, including histologically normal tissues. This setting enabled HiTIMED to apply its tumor-specific reference profile across all samples, generating low but non-zero tumor cell fractions in normal tissues. These inferred tumor-like signals may be biologically plausible and reflect early molecular alterations consistent with epigenetic field cancerization, in which adjacent or macroscopically normal tissues exhibit methylation changes associated with tumorigenesis. Alternatively, partial signal attribution may arise from technical noise, probe cross-reactivity, or shared epigenetic features between tumor and progenitor-like epithelial or stromal cells.

### Functional Annotation

Pathway membership for the VEGF signaling pathway was defined according to KEGG (KEGG ID: hsa04370)^28^, the PANTHER database^30^, and the Harmonizome database^29^. Hallmark gene sets, including the Hallmark Angiogenesis collection, were obtained from the Molecular Signatures Database^32^.

## Data Availability

All DNA methylation and gene expression data analyzed in this study were obtained from publicly available sources. Data are accessible through the respective repositories (https://portal.gdc.cancer.gov/, https://www.ncbi.nlm.nih.gov/geo/, and https://gtexportal.org/) under the terms provided by the original studies. Full accession details are provided in the Methods section. Any additional information or code used for data processing is available from the corresponding author upon reasonable request.

## Author Contributions

B.K. conceived and designed the study, performed the analyses, and wrote the manuscript with guidance and supervision from B.C.C. L.A.S. provided continuous feedback, discussion, and support with mathematical and statistical analyses, coding tools, and scientific content. Z.Z. contributed to coding support, data interpretation, and methodological discussions. H.L. provided expertise in breast cancer biology and additional coding assistance. All authors reviewed and approved the final manuscript.

