## Supplementary Tables for "Endothelial cell-specific DNA methylation alterations identified in breast cancer with epigenetic deconvolution"

| Discovery Set | | | Validation Set | | |
|---|---|---|---|---|---|
| **Characteristics** | **Normal tissue (n=340)** | **Tumor (n=609)** | **Characteristics** | **Normal tissue (n=75)** | **Tumor (n=471)** |
| **Age** | | | **Age** | | |
| Mean (sd) | 41 (13.5) | 57.5 (12.3) | Mean (sd) | 56.4 (14.8) | 56.4 (12.9) |
| Range | 13-82 | 29.5-92 | Range | 28-89 | 26-89 |
| **Tissue status, n (%)** | | | **Tissue status, n (%)** | | |
| Normal | 325 (96) | - | Normal | - | - |
| Tumor-adjacent normal | 15 (4) | - | Tumor-adjacent normal | 75 (100) | - |
| **Tumor Type, n (%)** | | | **Tumor Type, n (%)** | | |
| Ductal carcinoma in situ | - | 62 (10) | Ductal carcinoma in situ | - | 4 (0.8) |
| Invasive ductal carcinoma | | 547 (90) | Invasive ductal carcinoma | | 467 (99.2) |

**Table S1.** Study population and DNA methylation data

| Variable | Average VIF 1 | Average VIF 2 | Average VIF 3 |
|---|---|---|---|
| Age | 1.1 | 1.1 | 1.1 |
| Lymphocyte | 1.3 | 1.3 | 1.3 |
| Myeloid | 1.2 | 1.2 | 1.2 |
| Endothelial | 2.2 | 2.3 | 2.2 |
| Epithelial | 1.6 | 1.6 | 1.6 |
| Stromal | 2.2 | 2.3 | 2.2 |

**Table S2**. VIF estimates for model covariates

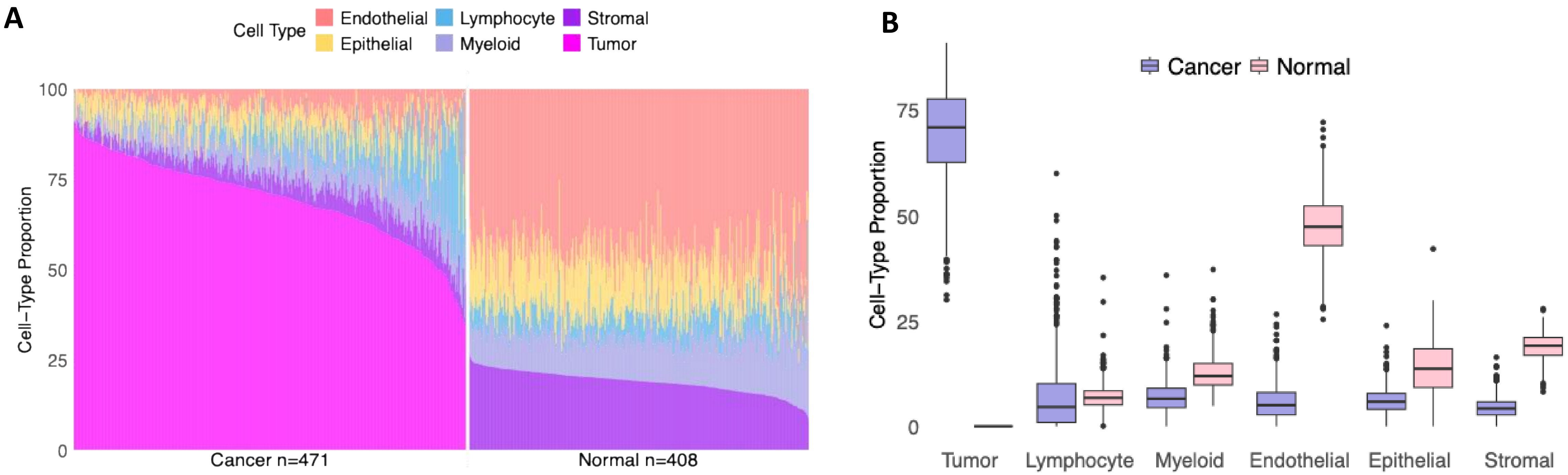


**Figure S1. Cell-type composition differences between breast cancer and normal breast tissue samples of the validation set**.
**A.** Stacked bar plots representing cell type proportions in validation set (breast cancer n=471, normal adjacent and normal breast n=408) ranked by tumor percentage in cancer tissue and stromal percentage in normal tissue. **B.** Boxplots show the estimated proportions of six major cell types—Tumor, Lymphocyte, Myeloid, Endothelial, Epithelial, and Stromal—in cancer (purple) and normal (pink) samples.

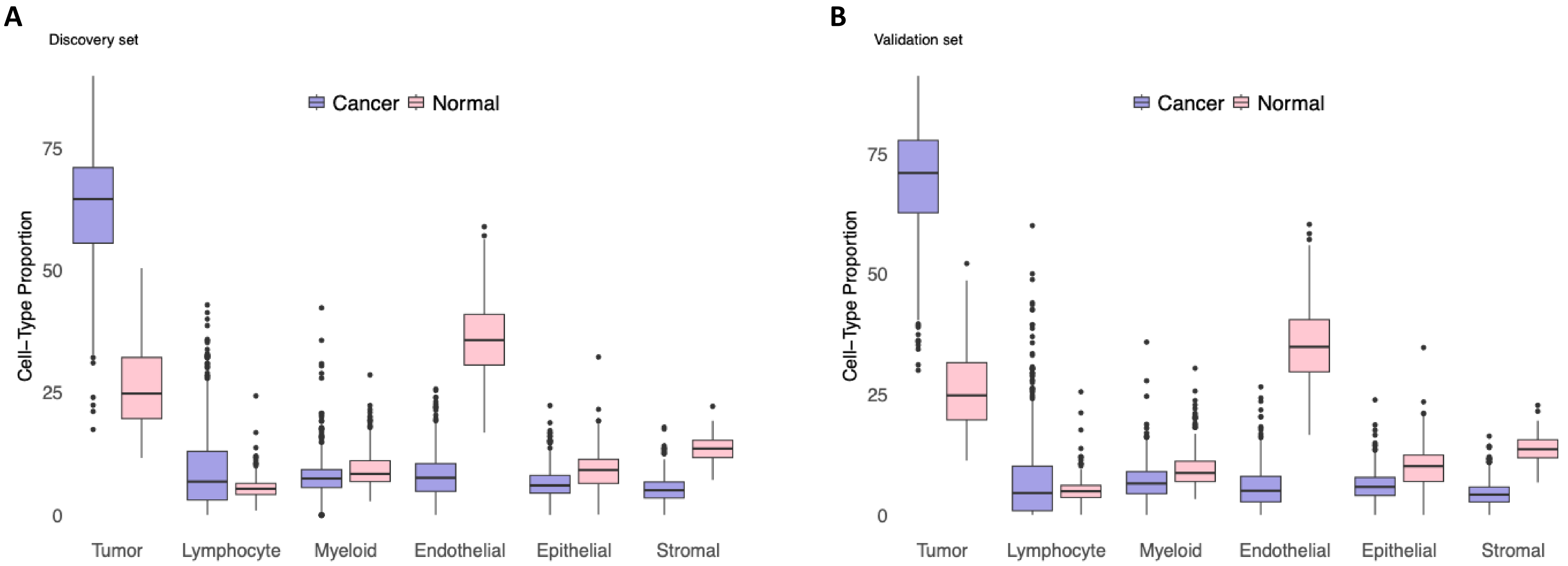


**Figure S2. Cancer and normal breast tissue cell-type proportions after re-estimating fractions with HiTIMED.**
Cell-type fractions were re-estimated by setting the tissue type to “tumor” for all samples for CellDMC analysis. **A.** Boxplots showing the estimated proportions of six major cell types — Tumor, Lymphocyte, Myeloid, Endothelial, Epithelial, and Stromal — in cancer (purple) and normal (pink) samples in the discovery dataset. **B.** Estimated cell-type proportions in the validation dataset.
